## Supplementary material for "Endomembrane targeting of human OAS1 p46 augments antiviral activity": Manuscript

**Table S1. Clinical and demographic information for COVID-19 and matched healthy control cohort.**

| Cohort | Number | M:F | Ancestry (self-reported) | rs10774671 genotype | | | Disease severity |
|---|---|---|---|---|---|---|---|
| | | | | AA | AG | GG | |
| COVID-19 Severe | 34 | 15:19 | 21% Caucasian<br>29% Latino<br>15% African American<br>6% Asian<br>15% American Indian<br>15% Other race/declined | 4<br>9<br>0<br>2<br>3<br>2 | 3<br>1<br>3<br>0<br>2<br>1 | 0<br>0<br>2<br>0<br>0<br>2 | Hospitalized, critical care unit, mechanical ventilation, or death |
| Ancestry matched controls | 99 | 47:52 | 34% Caucasian<br>22% Latino<br>19% African American<br>12% Asian<br>8% American Indian<br>3% Pacific Islander<br>2%Other race/declined | 16<br>10<br>6<br>4<br>1<br>1<br>0 | 14<br>11<br>9<br>8<br>4<br>1<br>1 | 4<br>1<br>4<br>0<br>2<br>1<br>1 | Not applicable |

**Table S2. Cells used in this study.**

| Name | Vendor/Source |
|---|---|
| *OAS1* KO HEK293T | Dan Stetson |
| *RNASEL* KO HEK293T | Dan Stetson |
| *OAS1* KO Huh7 | This manuscript |
| *OAS1* KO A549 | This manuscript |
| ACE2 HEK293T | This manuscript |
| *IRF3* KO 293FT | Dan Stetson |
| HEK293FT | ATCC |
| HEK293T | ATCC |
| A549 | ATCC |
| PH5CH8 | Michael Gale, Jr. |
| THP-1 | ATCC |
| Daudi | Saumen Sarkar |
| MDCK | Michael Gale, Jr. |
| Huh7 | Michael Gale, Jr. |
| Vero WHO | Michael Gale, Jr. |
| Vero R6 | Ralph Baric |
| PBMCs | Karen Cerosaletti, BRI |
| Human primary fibroblasts | Eric Allenspach, SCRI |

**Table S3. Antibodies used in this study.**

| Name | Vendor/Source | Catalog |
|---|---|---|
| OAS1 (D1W3A) Rabbit mAb | Cell Signaling Technology | 14498 |
| RNase L (D4B4J) Rabbit mAb | Cell Signaling Technology | 27281 |
| Golgin-97 (CDF4) Mouse mAb | Cell Signaling Technology | 97537 |
| Monoclonal Anti-PDIA3 antibody raised in mouse | Sigma-Aldrich | AMAB90988 |
| Golgin-97 (CDF4), Unconjugated, Species Reactivity: Human, Host: Mouse / IgG1 | Thermo Scientific | A-21270 |
| Monoclonal ANTI-FLAG® M2 antibody raised in mouse | Sigma-Aldrich | F3165 |
| J2 monoclonal antibody (mAb) anti dsRNA, mouse, IgG2a | Scicons | 10010200 |
| Mouse monoclonal antibody 9D5 anti dsRNA | Adam Geballe lab | N/A |
| β-Actin (13E5) Rabbit mAb | Cell Signaling Technology | 4970 |
| Goat anti Mouse IgG2a Alexa Fluor 647 | Thermo Fisher Scientific | A21241 |
| Goat anti Mouse IgG1 Alexa Fluor 594 | Thermo Fisher Scientific | A21125 |
| Goat anti Rabbit Alexa Fluor 488 | Thermo Fisher Scientific | A11008 |
| Zombie NIR fixable viability dye | BioLegend | 423105 |

**Table S4: Nucleic acids used in this study.**

| Primers | Sequence 5'-3' |
|---|---|
| OAS1 common For | GCTTGGTACCGAGCTCGGATCC |
| OAS1 common Rev | CAGCAGAATCCAGGAGCTCACTGG |
| OAS1 p42 For | AGTGAGCTCCTGGATTCTGCTGGTGAGACCTCCTGCTTCCTCCC |
| OAS1 p42 wt Rev | CTATAGAATAGGGCCCTCTAGATCAAGCTTCATGGAGAGGGGCAG |
| OAS1 p42 CaaX Rev | TATAGAATAGGGCCCTCTAGATCAGAGGATGGTGCAAGCTTCAT |
| OAS1 p46 For | CCCAGTGAGCTCCTGGATTCTGCTGGCTGAAAGCAACAGTGCAGACG |
| OAS1 p46 wt CaaX Rev | ACTATAGAATAGGGCCCTCTAGATCAGAGGATGGTGCAGGTCCA |
| OAS1 p46 mut CaaX Rev | ACTATAGAATAGGGCCCTCTAGATCAGAGGATGGTGGCGGTC |
| OAS1 p48 For | AGTGAGCTCCTGGATTCTGCTGACCCAGCACACTCCAGGCAG |
| OAS1 p46 C-term mut_F | TAAGAATTGGGATGGGTCCCCAG |
| OAS1 p46 C-term mut_R | GACACTATAGAATAGGGCCCTCTAGA |
| OAS1 p48 Rev | CTATAGAATAGGGCCCTCTAGATCAGGAGACCTGGGTTCTGTCC |
| OAS1DADA_F | GAAGACAACCAGGGCAGCGGCAGATCGGCCTCT |
| OAS1DADA_R | AGAGGCCGATCTGCCGCTGCCCTGGTT |
| OAS1FLAG_F | CGACTCACTATAGGGAGACCCAAGCTTGGTACCGAGCTCGATGGACTACAAAGAC |
| OAS1FLAG_R | GTCCAGAGATTTGGCTGGGGTATTTCTGAGATCCATCATGCTTGTCATCGTCATCCTTGTAATCGATG |
| EMCV_F | AGATCCGGATTGCCAGTCT |

| | |
|---|---|
| EMCV_R | CACTATCGTAGCCTTCACGTTG |
| EMCV_probe | /56-FAM/AT ATC GCA G/ZEN/G CTG GGT CCG T/3IABkFQ/ |
| IAV NP Fwd | CGTTCTCCATCAGTCTCCATC |
| IAV NP Rev | GAGTGACATCAAAATCATGGCG |
| OAS1 CFK mut SDM fwd | CTGAGGCCTGGCTGAATTACCCAGCCGCTGCGAATTGGGATGGGTCCCCAGTGA |
| OAS1 CFK mut SDM rev | TCACTGGGGACCCATCCCAATTCGCAGCGGCTGGGTAATTCAGCCAGGCCTCAG |
| OAS1 p46Δ32 SDM fwd | ACGATCCCAGGAGGTATCAGAAATGCACCATCCTC |
| OAS1 p46Δ32 SDM rev | GAGGATGGTGCATTTCTGATACCTCCTGGGATCGT |
| OAS1 p46Δ22 SDM fwd | ACATTGGAACACATGAGTACCCTTGCACCATCCTCT |
| OAS1 p46Δ22 SDM rev | AGAGGATGGTGCAAGGGTACTCATGTGTTCCAATGT |
| OAS1 p46Δ12 SDM fwd | CCCAGCACACTCCAGTGCACCATCCTCTGA |
| OAS1 p46Δ12 SDM rev | TCAGAGGATGGTGCACTGGAGTGTGCTGGG |
| OAS1 SDM common+CTIL fwd | AGAGGGGCAGGGATGAATGGCAGGGAGGACTAAAGAATCGTGCACAGCAGAATCCAGGAGCTCACTGGGGACC |
| OAS1 SDM common+CTIL rev | GGTCCCCAGTGAGCTCCTGGATTCTGCTGTGCACGATTCTTTAGTCCTCCCTGCCATTCATCCCTGCCCCTCT |
| OAS1 p46 Ala mut9 SDM fwd | CAGAGGATGGTGCAGGCCGCGTCCTCTTCTGCCTGT |
| OAS1 p46 Ala mut9 SDM rev | ACAGGCAGAAGAGGACGCGGCCTGCACCATCCTCTG |
| p52_F | CCCAGTGAGCTCCTGGATTCTGCTGCTGAAAGCAACAGTACAGACGATG |
| p52_R | ACTATAGAATAGGGCCCTCTAGATTATCTATGATAGGATAGAGGGCATAGAA |
| **S/AS oligos** | **Sequence 5'-3'** |
| OAS1 p42 WT S | GTGAGACCTCCTGCTTCCTCCCTGCCATTCATCCCTGCCCCTCTCCATGAAGCTTGA |
| OAS1 p42 WT AS | TCAAGCTTCATGGAGAGGGGCAGGGATGAATGGCAGGGAGGAAGCAGGAGGTCTCAC |
| OAS1 p42 CaaX S | GTGAGACCTCCTGCTTCCTCCCTGCCATTCATCCCTGCCCCTCTCCATGAAGCTTGCACCATCCTCTGA |
| OAS1 p42 CaaX AS | TCAGAGGATGGTGCAAGCTTCATGGAGAGGGGCAGGGATGAATGGCAGGGAGGAAGCAGGAGGTCTCAC |
| OAS1 p44 WT S | CCCAGTGAGCTCCTGGATTCTGCTGGTAAACCTCACACTGGTTGGCAGAAGGAACTATACCAATAATTAGTCTAGAGGGCCCTATTCTATAGTGT |
| OAS1 p44 WT AS | ACACTATAGAATAGGGCCCTCTAGACTAATTATTGGTATAGTTCCTTCTGCCAACCAGTGTGAGGTTTACCAGCAGAATCCAGGAGCTCACTGGG |
| **gBlocks (partial)** | **Sequence 5'-3'** |

| | |
|---|---|
| OAS1 p46 WT | GCTGAAAGCAACAGTGCAGACGATGAGACCGACGATCCCAGGAGGTATCAGAAATATGGTTACATTGGAACACATGAGTACCCTCATTTCTCTCATAGACCCAGCACACTCCAGGCAGCATCCACCCCACAGGCAGAAGAGGACTGGACCTGCACCATCCTCTGA |
| OAS1 p46 mutCaaX | GCTGAAAGCAACAGTGCAGACGATGAGACCGACGATCCCAGGAGGTATCAGAAATATGGTTACATTGGAACACATGAGTACCCTCATTTCTCTCATAGACCCAGCACACTCCAGGCAGCATCCACCCCACAGGCAGAAGAGGACTGGACCGCCACCATCCTCTGA |
| OAS1 p48 WT | ACCCAGCACACTCCAGGCAGCATCCACCCCACAGGCAGAAGAGGACTGGACCTGCACCATCCTCTGAATGCCAGTGCATCTTGGGGGAAAGGGCTCCAGTGTTATCTGGACCAGTTCCTTCATTTTCAGGTGGGACTCTTGATCCAGAGAGGACAAAGCTCCTCAGTGAGCTGGTGTATAATCCAGGACAGAACCCAGGTCTCCTGA |
| OAS1 p52 WT | CTGAAAGCAACAGTACAGACGATGAGACCGACGATCCCAGGACGTATCAGAAATATGGTTACATTGGAACACATGAGTACCCTCATTTCTCTCATAGACCCAGCACACTCCAGGCAGCATCCACCCCACAGGCAGAAGAGGACTGGACCTGCACCATCCTCTGAATGCCAGTGCATCTTGGGGGAAAGGGCTCCAGTGTTATCTGGACCAGTTCCTTCATTTTCAGGTGGGACTCTTGATCCAGAGAAGACAAAGCTCCTCAGTGAGCTGGTGTATAATCCAGGACAGAACCCAGGTCTCCTGACTCCTGGCCTTCTATGCCCTCTATCCTATCATAGATAA |
| OAS1 p46 Ala mut1 | TGCTTTAAGAATTGGGATGGGTCCCCAGTGAGCTCCTGGATTCTGCTGGCTGCCGCGGCAGCTGCAGACGATGAGACCGACGATCCCAGGAGGTATCAGAAATATGGTTACATTGGAACACATGAGTACCCTCATTTCTCTCATAGACCCAGCACACTCCAGGCAGCATCCACCCCACAGGCAGAAGAGGACTGGACCTGCACCATCCTCTGATCTAGAGGGCCCTATTCTATAGTGTCACCTA |
| OAS1 p46 Ala mut2 | TGCTTTAAGAATTGGGATGGGTCCCCAGTGAGCTCCTGGATTCTGCTGGCTGAAAGCAACAGTGCAGCGGCAGCTGCCGCAGCGCCCAGGAGGTATCAGAAATATGGTTACATTGGAACACATGAGTACCCTCATTTCTCTCATAGACCCAGCACACTCCAGGCAGCATCCACCCCACAGGCAGAAGAGGACTGGACCTGCACCATCCTCTGATCTAGAGGGCCCTATTCTATAGTGTCACCTA |
| OAS1 p46 Ala mut3 | TGCTTTAAGAATTGGGATGGGTCCCCAGTGAGCTCCTGGATTCTGCTGGCTGAAAGCAACAGTGCAGACGATGAGACCGACGATGCTGCCGCGGCAGCTGCATATGGTTACATTGGAACACATGAGTACCCTCATTTCTCTCATAGACCCAGCACACTCCAGGCAGCATCCACCCCACAGGCAGAAGAGGACTGGACCTGCACCATCCTCTGATCTAGAGGGCCCTATTCTATAGTGTCACCTA |
| OAS1 p46 Ala mut4 | TGCTTTAAGAATTGGGATGGGTCCCCAGTGAGCTCCTGGATTCTGCTGGCTGAAAGCAACAGTGCAGACGATGAGACCGACGATCCCAGGAGGTATCAGAAAGCTGCCGCGGCAGCTGCACATGAGTACCCTCATTTCTCTCATAGACCCAGCACACTCCAGGCAGCATCCACCCCACAGGCAGAAGAGGACTGGACCTGCACCATCCTCTGATCTAGAGGGCCCTATTCTATAGTGTCACCTA |
| OAS1 p46 Ala mut5 | TGCTTTAAGAATTGGGATGGGTCCCCAGTGAGCTCCTGGATTCTGCTGGCTGAAAGCAACAGTGCAGACGATGAGACCGACGATCCCAGGAGGTATCAGAAATATGGTTACATTGGAACAGCTGCCGCGGCAGCTGCATCTCATAGACCCAGCACACTCCAGGCAGCATCCACCCCACAGGCAGAAGAGGACTGGACCTGCACCATCCTCTGATCTAGAGGGCCCTATTCTATAGTGTCACCTA |
| OAS1 p46 Ala mut6 | TGCTTTAAGAATTGGGATGGGTCCCCAGTGAGCTCCTGGATTCTGCTGGCTGAAAGCAACAGTGCAGACGATGAGACCGACGATCCCAGGAGGTATCAGAAATATGGTTACATTGGAACACATGAGTACCCTCATTTCGCTGCCGCGGCAGCTGCACTCCAGGCAGCATCCACCCCACAGGCAGAAGAGGACTGGACCTGCACCATCCTCTGATCTAGAGGGCCCTATTCTATAGTGTCACCTA |

| | |
|---|---|
| OAS1 p46 Ala mut7 | TGCTTTAAGAATTGGGATGGGTCCCCAGTGAGCTCCTGGATTCTGCTGGC TGAAAGCAACAGTGCAGACGATGAGACCGACGATCCCAGGAGGTATCAG AAATATGGTTACATTGGAACACATGAGTACCCTCATTTCTCTCATAGACC CAGCACAGCTGCCGCGGCAGCTGCACCACAGGCAGAAGAGGACTGGACC TGCACCATCCTCTGATCTAGAGGGCCCTATTCTATAGTGTCACCTA |
| OAS1 p46 Ala mut8 | TGCTTTAAGAATTGGGATGGGTCCCCAGTGAGCTCCTGGATTCTGCTGGC TGAAAGCAACAGTGCAGACGATGAGACCGACGATCCCAGGAGGTATCAG AAATATGGTTACATTGGAACACATGAGTACCCTCATTTCTCTCATAGACC CAGCACACTCCAGGCAGCATCCACCGCTGCCGCGGCAGCTGCATGGACCT GCACCATCCTCTGATCTAGAGGGCCCTATTCTATAGTGTCACCTA |
| OAS1 p46 *Alligator mississippiensis* C-terminus | TGCTTTAAGAATTGGGATGGGTCCCCAGTGAGCTCCTGGATTCTGCTGCCT GAACAAACCCTCAAAGGGAGCAAGGGAGTCTGTGTTCGCTCTGTGGCTAA GCATGAAGCACGAAAACAGAAAGCTGCAGAACTTCAGCCTGTGCTGGTC TCTTCCTACAGCATCCCTGCCTTGGCCCCACAAGAGTTGGAAAAGCAACC CTCCTTTTGCAGCATACTCTGATCTAGAGGGCCCTATTCTATAGTGTCACC TA |
| OAS1 p46 *Bos taurus* C-terminus | TGCTTTAAGAATTGGGATGGGTCCCCAGTGAGCTCCTGGATTCTGCTGCC CCAAGAACACAGTGACCTGATGTTCCAGGCCTATGATTTTAGACAGCACT GTAGACCCTCTCCAGGAATCCAGTTCCACGGAGGAGCCTCTCCCCAGGTG GAAGAGAACTGGACATGTACCATCCTCTGATCTAGAGGGCCCTATTCTAT AGTGTCACCTA |
| OAS1 p46 *Pteropus alecto* C-terminus | TGCTTTAAGAATTGGGATGGGTCCCCAGTGAGCTCCTGGATTCTGCTGCC CTACGACACACCCCACGTGGAAGAGGACCAGTGGTGTGCCATCCTCTGAT CTAGAGGGCCCTATTCTATAGTGTCACCTA |
| OAS1 p46 *Vulpes vulpes* C-terminus | TGCTTTAAGAATTGGGATGGGTCCCCAGTGAGCTCCTGGATTCTGCTGCTT GAAGAAGACTATGAGGACAATTGGATAACCTCTGAACACAGGACATATT CATACCATGATTATGGTTGGCGCCCTGTATCCTCTGGGAGCCTCAACACA GGCATGACACAGTCCATTCCCCAGCAGGAAGAAAACTGGATGTGTACCAT CCTCTGATCTAGAGGGCCCTATTCTATAGTGTCACCTA |
| **siRNA** | **Sequence 5'-3'** |
| OAS1 | AGUCAAGCACUGGUACCAACAAUUUUGGUACCAGUG |
| p42 | GGUCACAAUCGAGGGUUAUUCCAGAAACCCUCGAU |
| p46 | AGAGAGAUUUAGAUAAGAUUCAUUCUCUUAUCUAAA |

**Table S5. Viruses used in this study.**

| Name | Vendor/Source | Catalog |
|---|---|---|
| Encephalomyocarditis virus | ATCC | VR-1762 |
| West Nile virus Texas | Michael Gale, Jr. | |
| CVB3-Nancy | Raul Andino | |
| Influenza virus A/PR/8/34 | ATCC | |
| Influenza A virus A/Udorn/72 H3N2 R38A | Michael Gale, Jr. | |
| Indiana vesiculovirus (VSV-GFP) | Michael Gale, Jr. | |
| SARS-CoV-2 strain USA/WA-1/2020 | Michael Gale, Jr. | |
| ZIKV MR766 | Michael Gale, Jr. | |
